# Genomic Signatures of Fine-Scale Local Adaptation in Atlantic Salmon Suggest Involvement of Sexual Maturation, Energy Homeostasis, Behaviour, and Immune Defence-Related Genes

**DOI:** 10.1101/271528

**Authors:** Victoria L. Pritchard, Hannu Mäkinen, Juha-Pekka Vähä, Jaakko Erkinaro, Panu Orell, Craig R. Primmer

**Author notes:** Present address: Association for Water and Environment of Western Uusimaa, POB 51, FI-08101 Lohja, Finland.

## Abstract

Elucidating the genetic basis of adaptation to the local environment can improve our understanding of how the diversity of life has evolved. In this study we used a dense SNP array to identify candidate loci underlying fine-scale local adaptation within a large Atlantic salmon (*Salmo salar*) population. By combining outlier, gene-environment association, and haplotype homozygosity analyses, we identified multiple regions of the genome with strong evidence for diversifying selection. Several of these candidate regions had previously been identified in other studies, demonstrating that the same loci be adaptively important in Atlantic salmon at sub-drainage, regional and continental scales. Notably, we identified signals consistent with local selection around genes associated with variation in sexual maturation, energy homeostasis, behaviour, and immune defence. These included the large-effect age-at-matunty gene *vgll3*, the known obesity gene *mc4r*, and major histocompatibility complex II. Most strikingly, we confirmed a genomic region on Ssa09 that was extremely differentiated among subpopulations, and that is also a candidate for local selection over the global range of Atlantic salmon. This region co-localized with a genomic region strongly associated with spawning ecotype in sockeye salmon (*Oncorhynchus nerka*), with circumstantial evidence that the same gene (*six6*) may be the selective target in both cases. The phenotypic effect of this region in Atlantic salmon remains cryptic, although allelic variation is related to river flow volume and co-vanes with timing of the return spawning migration. Our results further inform management of Atlantic salmon and open multiple avenues for future research.

## INTRODUCTION

Understanding how the diversity of life on earth has evolved is one of the central questions in biology. Fundamental to this is elucidating how change at the genomic level underlies phenotypic change as populations adapt to their local environment, and how this change ultimately contributes to evolutionary radiations. Central questions include the relative roles of polygenic variation vs. genes of large effect (Yeaman & Whitlock, 2011), the importance of protein coding vs. gene expression variation (Fraser, 2013), the contribution of chromosome rearrangements (Kirkpatrick & Barton, 2006) and whether parallel, independently evolved adaptations have the same genetic basis across populations (Conte, Arnegard, Peichel, & Schluter, 2012). Resolving such questions contributes to the ultimate aim of linking genotype, phenotype and fitness (Barrett et al. 2011). One limitation until recently has been the lack of dense marker panels and annotated genomes for non-model species, which impedes identification of the loci driving statistical signals of selection (Vilas, Pérez-Figueroa, and Caballero 2012).

Salmonid fishes, which include charr, salmon, trout and whitefish, are splendid models for the study of local adaptation and diversification. Strong philopatry in breeding location and/or separation into different water bodies means that fine-scale genetic differentiation is a characteristic of salmonid populations (Fraser, Weir, Bernatchez, Hansen, & Taylor, 2011). Both allopatric and sympatric populations can express a wide range of phenotypic diversity, including variation in age at maturity, migratory strategy, breeding time and location, and exploitation of different ecological niches (e.g. Bernatchez et al., 2010; Dodson, Aubin-Horth, Thériault, & Páez, 2013; Jonsson & Jonsson, 2001; Quinn, McGinnity, Reed, & Bradford, 2016). Evolutionary diversification may have been facilitated by the whole genome duplication that occurred at the base of the salmonid radiation (Lien et al., 2016). Much of this diversity is heritable (Carlson & Seamons, 2008), and multiple studies have found reduced fitness of salmonids in non-local environments (Fraser et al., 2011; O’Toole et al., 2015). While many fitness-associated traits in salmonids have been well characterized at the quantitative genetic level, most of the underlying loci are yet to be identified (Fraser, Weir, Bernatchez, Hansen, & Taylor, 2011; Garcia de Leaniz et al., 2007).

New genomic tools are enabling us to unpick the molecular genetic basis of local adaptation in salmonids (Elmer, 2016). This can both improve our understanding of evolutionary diversification in general and help guide management of these ecologically and culturally important taxa (e.g. Prince et al., 2017). Recent studies have identified suites of loci potentially involved with adaptation to the natal or migratory environment (e.g. Micheletti, Matala, Matala, & Narum, 2017; Moore et al., 2017), associated with ecotypic differentiation (e.g. Veale & Russello, 2017a), or responding to selection (e.g. Liu et al. 2017). One finding of such studies is that life history diversification, in particular, can be underlain by small genomic changes of large effect. For example, Atlantic salmon (*Salmo salar*) vary in their age at maturity, with some individuals returning to freshwater to spawn after one year feeding at sea (‘one-seawinter fish’), and others spending longer in the ocean and returning at a much larger size (‘multi-seawinter fish’). Barson et al. (2015) showed that a single locus containing the gene *vgll3* explained nearly 40% of this variation in age at maturity across three evolutionary lineages in Europe (see also Ayllon et al., 2015). This locus appears also to be associated with sea age at maturity in North American Atlantic salmon (Kusche et al., 2017), and is additionally located within a QTL underlying early (‘precocious’) freshwater maturation (Lepais, Manicki, Glise, Buoro, & Bardonnet, 2017). The choice between residency and anadromy in many populations of rainbow trout/ steelhead (*Oncorhynchus mykiss*) is associated with a genomic region on chromosome Omy05 (Hecht, Campbell, Holecek, & Narum, 2013; Hecht, Thrower, Hale, Miller, & Nichols, 2012; Martínez, Garza, & Pearse, 2011; Nichols, Edo, Wheeler, & Thorgaard, 2008; Pearse, Miller, Abadía-Cardoso, & Garza, 2014). Sockeye salmon (*O. nerka*) exhibit ecotypic variation in spawning site, with both anadromous and resident forms spawning in streams(/rivers) or lake shores(/benthos): different spawning ecotypes can co-occur within the same populations (Larson et al., 2017; Nichols, Kozfkay, & Narum, 2016; Veale & Russello, 2017b). Veale & Russello (2017a) showed that this ecotypic differentiation is associated with two strongly diverged haplotypes encompassing the *lrrc9* gene. This relationship held throughout the species’ range: in some populations, it was almost entirely Mendelian, with one homozygote determining shore spawning. Finally, in one case, independently arising mutational variation at the same locus underlies an ecologically important life history trait in two species: variation in the *greb1 l* gene is associated with premature spawning migration in both steelhead and chinook salmon (*O. tshawytscha*) (Hess, Zendt, Matala, & Narum, 2016; Prince et al., 2017)

One of the world’s largest surviving wild Atlantic salmon stocks reproduces in the Teno River (Norwegian: Tana; Sámi: Deatnu, Fig. 1) of subarctic Finland and Norway. The Teno drains a catchment of 16,386km^2^ and includes more than 30 different-sized tributaries accessible to adult salmon: up to 100,000 individuals return to spawn each year. Previous studies have demonstrated significant, temporally stable, population genetic substructure within the Teno stock corresponding to different spawning locations (Vähä, Erkinaro, Falkegård, Orell, & Niemelä, 2017; Vähä, Erkinaro, Niemelä, & Primmer, 2008; global F_st_ from microsatellites ≈ 0.065). This genetic structure is higher than that observed across multiple rivers in other parts of the Atlantic salmon range (Vähä, Erkinaro, Falkegård, Orell, & Niemelä, 2017). Thus, there is strong potential for differential local adaptation within the system. Teno salmon subpopulations vary in the timing of their re-entry to freshwater to spawn (‘run timing’; June-August; Vähä et al., 2011), and the proportion of multi-seawinter adults among spawners (Vähä, Erkinaro, Niemelä, & Primmer, 2007). Recently, Aykanat et al. (2015) demonstrated that adult salmon captured in the Teno mainstem belong to two genetically distinct subpopulations that differ in their freshwater and marine growth rate. Otherwise, no detailed studies of phenotypic differentiation among subpopulations have been performed.

**Figure 1:**
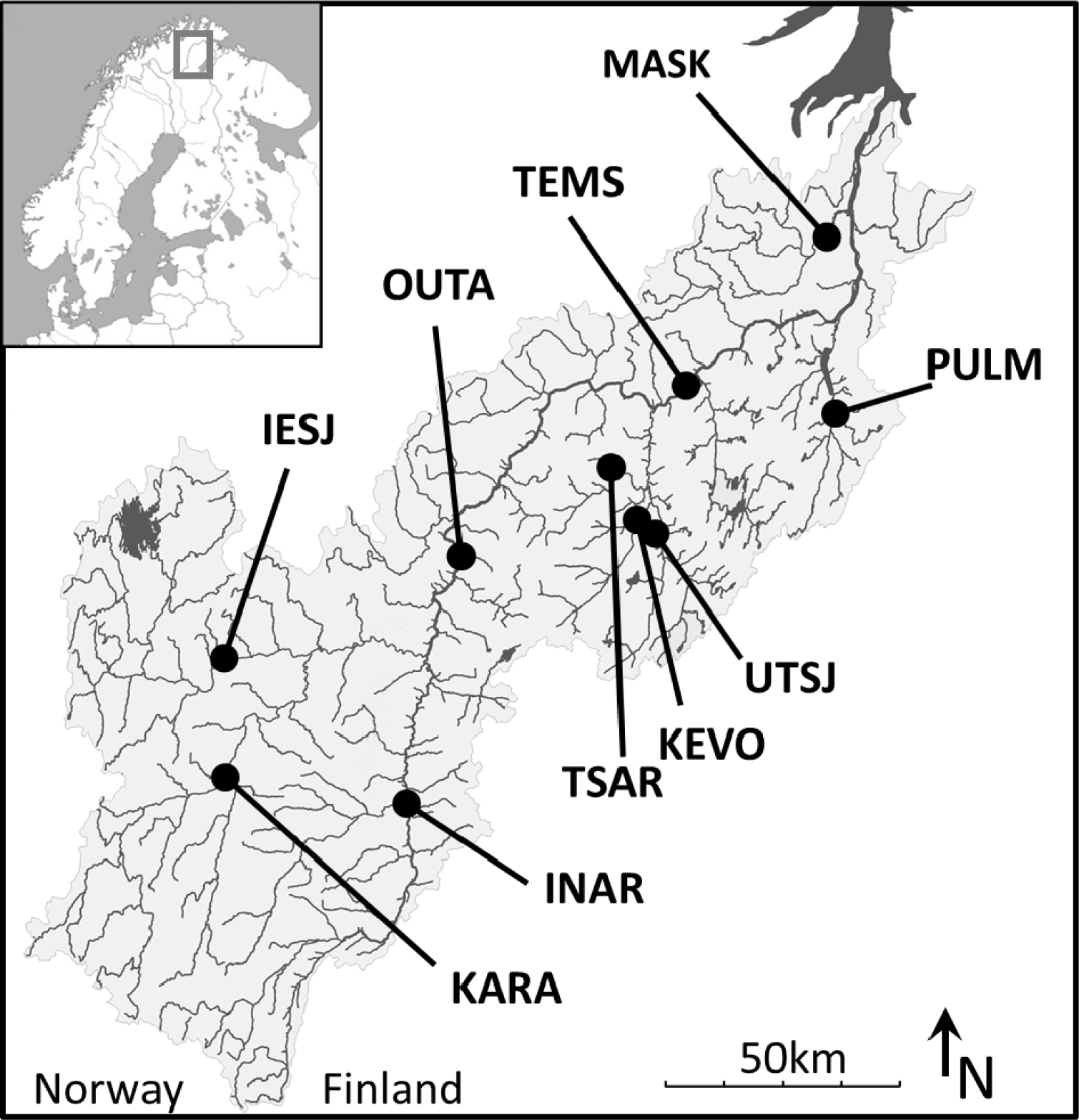
Location of juvenile sampling sites

In this study, we used a dense marker panel (198,829 SNPs) to examine genomic variation among Teno Atlantic salmon subpopulations, with the aim of identifying regions of the genome that potentially contribute to fine-scale local adaptation. To examine whether these candidate regions might also be under differential selection at broader geographic scales, we compared our results to previous studies of Atlantic salmon throughout their global range. We also examined our candidate regions for any of the large-effect genes implicated in life history diversification in other salmonid species.

## MATERIALS AND METHODS

### Sample Collection, Preparation and Genotyping

To ensure that samples represented the local breeding subpopulation, we genotyped juvenile salmon collected before their ocean migration (age 0 and 1). Sample collection and DNA extraction is described in Vähä et al. (2017). To minimize sampling of siblings, electrofishing sites were separated by 50-100m and only a single individual of each age was collected at each site. We selected 10 locations in the Teno known to harbour genetically distinct subpopulations (Aykanat et al., 2015; Vähä et al., 2017, 2008) and which represent a range of environmental variation (Vähä et al., 2007): Teno mainstem: Garnjarga (TEMS, n=24), Teno mainstem: Outakoski (OUTA, n=24), Inarijoki (1NAR, n=21), lower Iešjohka (IESJ, n=21), Maskejohka (MASK, n=20), Kárásjohka (KARA, n=22), Tsarsjoki/Carsejohka (TSARS, n=21), Utsjoki/Ohcejohka (UTSJ, n=21), Kevojoki/Geavvujohka (KEVO, n=21), Yla Pulmankijarvi/Buolbmátjohka (PULM, n=20) (Fig. 1, Table 1). We assessed concentration and degradation of the previously-extracted DNA using a Nanodrop ND-1000 spectrophotometer (Thermo Fisher Scientific Inc.) and by agarose gel visualization, and performed re-extractions where necessary. DNA of sufficient quality was standardized to a Nanodrop-estimated concentration of 15ng/ul and sent to the Center for Integrative Genetics (CIGENE), Ås, Norway for genotyping.

**Table 1:** Sample site environmental and phenotypic variables. Median run time is estimated from 1SW adults only (Vähä et al., 2011). Percentage of multi-scawinter females is rounded to the nearest 5%.

| SITE | Latitude | Longitude | Catch-<br>ment<br>(km<sup>2</sup>) | Distance<br>(km) | Elevation<br>(m) | Access-<br>ibility | % MSW<br>females | Median<br>run time<br>(day) |
| --- | --- | --- | --- | --- | --- | --- | --- | --- |
| IESJ | 69° 24' 23.49" | 24° 40' 50.93" | 1900 | 259 | 230 | 4 | 95 | 185 |
| INAR | 69° 8' 13.67" | 25° 44' 33.39" | 1257 | 245 | 159 | 4 | 45 | 185 |
| KARA | 69° 10' 28.08" | 24° 41' 39.30" | 1235 | 285 | 244 | 4 | 90 | 184 |
| KEVO | 69° 42' 11.52" | 26° 57' 27.49" | 494 | 127 | 94 | 5 | 10 | 178 |
| MASK | 70° 12' 43.50" | 27° 56' 36.71" | 463 | 41 | 56 | 2 | 60 | 187.5 |
| OUTA | 69° 38' 0.45" | 25° 57' 41.63" | 8387 | 175 | 115 | 4 | 45 | 185 |
| PULM | 69° 53' 36.56" | 28° 1' 29.77" | 598 | 76 | 22 | 2 | 20 | 172 |
| TEMS | 69° 55' 54.79" | 27° 8' 58.64" | 11035 | 93 | 60 | 3 | 75 | 190 |
| TSAR | 69° 46' 58.03" | 26° 54' 25.39" | 236 | 125 | 170 | 6 | 20 | 173 |
| UTSJ | 69° 39' 51.21" | 27° 4' 16.28" | 441 | 131 | 107 | 4 | 85 | 184.5 |

Samples were genotyped for 220,000 SNPs using a custom Affymetrix Axiom SNP array on a GeneTitan genotyping platform, as in Barson et al. (2015). SNPs on this 220K array have known locations on the NCBI RefSeq Atlantic salmon genome (Lien et al., 2016, http://www.ncbi.nkn.nili.gov/genome/annotation_euk/Salmo_salar/100/). To ensure correct identification of genotype clusters, we applied the Affymetrix Best Practices Protocol for SNP calling simultaneously to our samples and ≈1800 individuals genotyped in previous studies (Barson et al., 2015; Pritchard et al., 2016).

For quality control and analyses we also used two pre-existing 220K SNP datasets: a dataset of known aquaculture escapees (n=192, Pritchard et al., 2016), and a dataset of adults collected from the Teno mainstem (n=463) and the neighbouring river Borselva (n=17) (Barson et al., 2015). The adult Teno mainstem sample included the individuals genotyped with a 7K SNP array by Johnston et al. (2014) and Aykanat et al. (2015).

### Genotyping quality control

Two hundred and two out of 215 samples passed Affymetrix quality thresholds. Subsequent quality control was performed using PLINK v.1.90 (Chang et al., 2015; Purcell et al., 2007). First, we removed 1,112 SNPs not mapped to an assembled *S. salar* chromosome and 35 SNPs with known off-target variants. We then used the dataset of adults collected from the Teno mainstem to identify SNPs that deviated from Hardy-Weinberg equilibrium at p<0.001 in either of the two subpopulations of Aykanat et al. (2015; Pop1: n=303; Pop2: n=121). We found 2,750 such SNPs; we considered these to have technical genotyping problems and removed them. To identify full sibs within our juvenile samples, we split them by subpopulation, excluded SNPs with >2% missing data or MAF<0.05 within each subpopulation, performed a linkage disequilibrium (LD) pruning step (P LINK command: *--indep 50 5 1.4*), used the *--genome* function to estimate genome-wide identity-by-descent between each pair of individuals, and graphically explored the results using ggplot2 in R 3.1.2 (R Core Team, 2015; Wickham, 2009). To identify juveniles with possible domestic parentage we added the dataset of aquaculture escapees, performed SNP filtering and LD pruning as described above, applied a two-dimensional multidimensional scaling analysis (MDS) to the genome-wide identity-by-state (IBS) matrix in PLINK, and graphically visualized the results. Following removal of 13 putative sibs and one individual with possible escapee ancestry, we excluded all SNPs with a MAF < 0.05 or >10% missing genotypes. The final dataset included 198,829 SNPs and 188 individuals (TEMS, n= 24; OUTA, n=22; PULM, n=13; IESJ n=21; INAR n=20; KARA n=19; TSAR n=21; UTSJ n=21; KEVO n=19; MASK n=8). Proportion of missing genotypes per individual ranged from 0.07% to 3.24% (median 0.20%).

### Population genomic characteristics

We calculated subpopulation H_e_ from genome-wide minor allele frequencies returned by PLINK We also used PLINK to estimate unbiased pairwise F_st_ (Weir & Cockerham, 1984). We explored subpopulation structure among the juveniles by performing an MDS analysis on the LD-pruned final dataset, as described above. We investigated how juvenile samples related to the adult samples from the Teno mainstem by combining the two datasets and performing the same analysis. We assigned adults to the two mainstem subpopulations of Aykanat et al. (2015) using the Structure results from that study (available: https://doi.org/10.5061/dryad.7t4n0), with a cut-off of q=0.9 for subpopulation membership.

### Enrironmental variation

We investigated four broad-scale environmental characteristics previously shown to influence population structure in the Teno (Vähä, Erkinaro, Niemelä, & Primmer, 2007; Table 1). We estimated waterway distance of the sampling site from the Teno mouth using the package ‘riverdist’ in R (https://cran.r-project.org/web/packages/riverdist/index.html). We obtained altitude using Google maps. We estimated upstream catchment area for each site (a surrogate for flow volume) by summing sub-basin catchment areas obtained from the Finnish Center for Economic Development, Transport and the Environment. We based presence of downstream barriers on data for nearby sites in Vähä, Erkinaro, Niemelä, & Primmer (2007). As these four variables are non-orthologous, we converted them into principle components: standardized values were provided to the function *prcomp* in R with default parameters.

### Phenotypic variation

To explore the relationship between these environmental variables and known phenotypic variation within the Teno, we extracted median run time and proportion of multi-seawinter females for each sampled subpopulation from Vähä et al. (2011) (Table 1) and examined their relationship to the principal components using a Kendall rank correlation test in R.

### Identification of candidate genome regions responding to local selection

Best practice requires a combination of approaches to identify candidate genome regions responding to local selection (De Villemereuil, Frichot, Bazin, François, & Gaggiotti, 2014). We therefore a) identified markers unusually highly differentiated among subpopulations, based on F_st_ or equivalent statistics; b) examined association of allele frequencies with environmental parameters; and c) examined patterns of haplotype homozygosity within subpopulations, indicative of selective sweeps on particular alleles. We used PLINK or PGD Spider (Lischer & Excoffier, 2012) to convert among input files. We used Beagle 4.1 (Browning & Browning, 2007) with the complete juvenile dataset to impute missing genotypes or infer phasing where necessary, To assess whether several candidate SNPs could be labelling a single selected locus, we identified sets of neighbouring SNPs in high LD (‘haploblocks’) using *--blocks* in PLINK. Because default parameters in PLINK often returned multiple small haploblocks that appeared to break up a single selective sweep, we used relaxed parameters (*--no-small-max-span --blocks-inform-frac 0.8 --blocks-max-kb 5000 --blocks-strong-lowci 0.55 --blocks-strong-highci 0.85 --blocks-recomb-highci 0.8*). We defined haploblock bounds as the positions halfway between the outermost haploblock SNPs and their closest non-haploblock SNPs. We condensed any abutting haploblocks containing candidate SNPs and within 10kB of each other into a single block.

#### Outlier approaches

we used three approaches to identify markers with high among-population allelic variation. First, we estimated among-population F_st_ for each marker using OutFlank (Whitlock & Lotterhos, 2015). Second, we used Bayescan (Foll & Gaggiotti, 2008), which assumes an island model of migration, with N_e_ and migration rate allowed to vary among subpopulations; diversifying selection is indicated by positive values of alpha, the locus-specific component of F_st_. We specified *–pr-odds* 100, kept other parameters default, and assessed convergence over three replicate runs. Third, we used BayEnv2 (Coop, Witonsky, Di Rienzo, & Pritchard, 2010; Gunther & Coop, 2013), which accounts for population structure and unequal sampling by estimating a variance - covariance matrix of allele frequencies across populations, and returns the *X^T^X* statistic, which is a measure of the deviation of each locus from the underlying matrix and hence the likelihood that the locus is under diversifying selection. We estimated the population covariance matrix from an LD pruned subset of 33,133 SNPs (PLINK *--indep 50 5 1.4*), using 200,000 iterations. We supplied this matrix in three replicate runs of BayEnv2 for each SNP (100,000 iterations), and took the median of the three *X*^*T*^*X* scores.

#### Association with environmental variables

Two principle components explained 82% of the total environmental variation and were included in the analysis. We used two approaches to identify markers associated with PC1 or PC2. BayEnv2 investigates allele frequency-environment associations accounting for population structure and sampling variance as described above. We used the previous subpopulation variance-covariance matrix, performed 200,000 iterations, and ran the analysis five times. As we observed little concordance between Bayes Factor rankings of SNPs among replicate runs, but much better concordance between rankings based on Spearman’s *P*, we used the latter as our informative measure. Such Bayes Factor discordance among replicate runs been noted elsewhere, and may not be solved by further increasing run length (Blair, Granka, & Feldman, 2014). We used the median of the absolute values of *P* over the five runs as our test statistic. Latent factor mixed models (LFMM, Frichot, Schoville, Bouchard, & François, 2013) investigate allele frequency-environment associations using mixed models in which latent variables account for population structure. For the LFMM analyses we imputed missing genotypes, specified 10 latent factors, performed 10,000 iterations with 5,000 burn-in, ran the analysis five times, and took the median z-score. We confirmed that 10 was a suitable number of factors using PCA with a Tracy-Widom test in the R package LEA (Frichot, Mathieu, Trouillon, Bouchard, & François, 2014).

#### Haplotype homozygosity

We used two approaches to examine subpopulations for elevated haplotype homozygosity, reflecting a selective sweep on an allele within that haplotype. The cross-population extended haplotype homozygosity (EHH) test (XP-EHH, Sabeti et al., 2007) compares EHH at the same site between two populations and thus can be used to identify selective sweeps that have occurred in one population but not in the other. We estimated normalized XP-EHH for each SNP using `selscan` (Szpiech & Hernandez, 2014; *--max-gap* 2Mb, all other parameters default). We selected the Teno mainstem (TEMS) subpopulation as a reference and compared each of the 9 other subpopulations; thus, negative XP-EHH values indicate higher haplotype homozygosity in the TEMS subpopulation and positive values vice-versa. We normalized XP-EHH across all chromosomes within each pairwise comparison. To obtain an overall score for each SNP we took the maximum absolute normalized score over all comparisons.

HapFLK (Fariello, Boitard, Naya, SanCristobal, & Servin, 2013) organizes markers into haplotype clusters using a multipoint linkage disequilibrium model and then measures haplotype frequency differentiation between populations, accounting for population structure using a population tree. For the HapFLK analysis, we included Borselva as the outgroup and inferred the population tree from the same 33,133 SNPs used to estimate the variance-covariance matrix for BayEnv2. We ran HapFLK for each chromosome separately, specifying 10 or 15 haplotype clusters (*K*) and 10 EM runs (*nfit*). The same outlying regions were identified independent of K, and we retained the HapFLK scores for each SNP from the K=10 analysis.

#### Significance testing and combined evidence

The significance of observed outliers or environmental associations in genome scans is frequently assessed by comparing observed scores to the expected distribution of scores in the absence of selection. However, modelling this neutral distribution requires assumptions about the demographic history of the populations under study that are rarely met. Correspondingly, simulation studies have demonstrated varying levels of Type I and Type II error depending on the true underlying scenario and the sampling scheme (Lotterhos & Whitlock, 2014; Narum & Hess, 2011; De Villemereuil et al., 2014). In a similar vein, the statistical properties of test scores used to investigate haplotype homozygosity are not well characterized (Vatsiou, Bazin, & Gaggiotti, 2015). We did not have a way to a-priori identify a set of selectively neutral SNPs (Lotterhos & Whitlock, 2014) from which we could obtain an expected distribution of scores. Further, neutral or non-locally-adaptive population genetic processes such as allele surfing and purifying selection can generate high test scores, meaning that apparently ‘significant’ loci still require validation by further studies (Charlesworth, Nordborg, & Charlesworth, 1997; Edmonds, Lillie, & Cavalli-Sforza, 2004). Given these considerations, we selected our loci of interest based on the empirical distribution of test scores. For each test, we ranked SNPs by test score and retained the top-ranked 0.5% (equivalent to empirical p<0.005 considering all 198,829 SNPs). We combined evidence over these nine sets of SNPs to identify ‘candidate SNPs’ as follows:

i. SNPs in the top 0.5% set in both the LFMM and BayEnv2 analyses were considered environmentally associated SNPs (‘PC1’ or ‘PC2’ SNPs).
ii. SNPs in the top set in either the LFMM or the BayEnv2 analysis, and also in the top set in at least two of three outlier analyses were also considered environmentally associated SNPs (‘PC1’ or ‘PC2’ SNPs).
iii. SNPs in the top set in at least two of the three outlier analyses, but not in the top set of either environmental analysis, were considered outlier SNPs (‘OUT’ SNPs).
iv. SNPs in the top set in both the XP-EHH and HapFLK analysis, but not in other analyses, were considered extended haplotype homozygosity SNPs (‘EHH’ SNPs).
v. We identified a subset of haploblocks that were particularly strong candidates to contain loci under diversifying selection by combining results from the outlier/environmental analyses with results from the XP-EHH and HapFLK analyses. ‘Candidate haploblocks’ contained at least one PC1, PC2 or OUT SNP and at least one SNP in the top 0.5% set of either the XP-EHH or HapFLK analysis.

#### Additional analyses

on the basis of our results, we performed an additional, exploratory, LFMM analysis using median run time as the dependent variable, following the method described above.

### Annotation of candidate regions

We found genes associated with candididate SNPs and haploblocks, using information from NCBI *Salmo salar* Annotation Release 100 provided in the R package Ssa.RefSeq.db (Grammes, 2016). We excluded non-coding RNAs and pseudogenes from the annotation, and converted gene, SNP and haploblock positions into bed format. To annotate candidate SNPs (PC1, PC2, OUT or EHH), we first used the bedtools function *intersect* to find overlapping genes (Quinlan & Hall, 2010). For SNPs in intergenic regions, we then used the function *closest* to find the nearest downstream gene, assuming that a selected variant could be in a 5’ regulatory region. We also identified the intersecting or closest downstream gene for each of the 198,829 SNPs to create the background gene set for a GO term enrichment analysis (see below). For candidate haploblocks, we annotated overlapping and closest downstream genes.

### GO term enrichment analysis

We obtained gene ontology (GO) terms for each annotated *S. salar* gene from Ssa.RefSeq.db, and performed GO term enrichment analyses using the R package topGO 2.28.0 (Alexa & Rahnenfuhrer, 2010). We assessed enrichment compared to the background described above using the Fisher exact test with the *weight01* algorithm. We performed four analyses: i) genes associated with all candidate SNPs; ii) genes associated with PC1 SNPs; iii) genes associated with PC2 SNPs; iv) genes associated with OUT SNPs. Where a gene was associated with both a PC1/PC2 and an OUT SNP, we included it in the PC1 or PC2 group only. We did not apply a multiple-testing correction as the level at which this should be performed for GO term enrichment analyses is unclear (Alexa & Rahnenfuhrer, 2010); instead we examined top-ranking GO terms in each test.

### Comparison of outliers to previous results

Several previous studies have used a 7K SNP array to identify regions of the genome putatively under differential selection among *S. salar* populations (Bourret, Kent, et al., 2013; Bourret, Dionne, Kent, Lien, & Bernatchez, 2013; Gutierrez, Yáñez, & Davidson, 2016; Jeffery et al., 2017; Liu et al., 2017; Mäkinen, Vasemägi, McGinnity, Cross, & Primmer, 2015; Moore et al., 2014; Perrier, Bourret, Kent, & Bernatchez, 2013); discriminating populations (Karlsson, Moen, Lien, Glover, & Hindar, 2011) or associated with phenotypic traits (Gutierrez, Yáñez, Fukui, Swift, & Davidson, 2015; Johnston et al., 2014). To compare our results with the location of SNPs of interest identified in these studies, we obtained flanking sequences for the 7K SNPs from NCBI dbSNP (www.ncbi.nlm.nih.gov/projects/SNP/) and aligned them with the *S. salar* genome using bwa-mem with default parameters (Li, 2013; Li & Durbin, 2009). We only retained unambiguously mapped sequences with MQ ≥ 60. We considered a SNP to map to one of our candidate regions if it was within the haploblock boundary.

To further explore one strong candidate region for local selection, we aligned two *O. nerka* RAD-tag sequences to the annotated *S. salar* and/or *O. mykiss* genomes using bwa-mem (Omyk_1.0 available: https://www.ncbi.nlm.nih.gov/assembly/GCF_002163495.1). RAD-tag 68810/57884 was associated with spawning ecotype by Veale & Russello (2017b) and Nichols, Kozfkay, & Narum (2016), and maps to the *lrrc9* gene (Veale & Russello 2017a). RAD-tag 24343/64477/41305 was associated with spawning ecotype by Veale & Russello (2017b), Nichols, Kozfkay, & Narum (2016), and Larson et al. (2017), but not previously annotated. Finally, we used bwa-mem to align the zebrafish variant of the *six6* retinal enhancer from Conte et al. (2010) to the *S. salar* genome.

## RESULTS

### linrironmental variation and population structure

PC1 explained 58% of the variation observed within the environmental data and was related to the correlated variables distance (loading: −0.58), elevation (−0.63) and accessibility (−.47); thus we considered it to broadly represent adult migration distance. PC2 explained a further 24% of variation and was strongly related to upstream catchment area (loading: 0.96); thus we considered it to represent river flow volume (Table S1). Position of sampling sites along the two environmental PCs is shown in Fig. S1. Median runtime was positively related to PC2 (Fig. S2).

Pairwise F_st_ between samples is shown in Table S2; global F_st_ was 0.067, similar to previous microsatellite-based analyses (Vähä et al. 2017). We observed slightly lower genome-wide heterozygosity in the tributary subpopulations (H_e_, KEVO: 0.338; MASK: 0.349; PULM: 0.320; TSAR: 0.294; UTSJ: 0.343) than in the mainstem and headwater subpopulations (IESJ: 0.370; INAR: 0.362; KARA: 0.363; OUTA 0.372; TEMS: 0.373). Correspondingly, MDS visualization of population structure revealed differentiation at two hierarchical levels. Analysis of the entire dataset showed the tributary subpopulations KEVO, TSAR, UTSJ and PULM to be strongly differentiated from one another and the mainstem, headwater and MASK subpopulations (Fig. 2). Excluding these tributary subpopulations and repeating the analysis also confirmed genomic divergence among the remaining six subpopulations (Fig. 2), although at least two individuals from OUTA were possible migrants from other sites (Fig. 2).

Comparing Teno mainstem and headwater juveniles to adults caught in the mainstem revealed, clearly, that the two genetically differentiated subpopulations described in Aykanat et al. (2015) derived from different spawning locations (Fig S3). ‘Subpopulation 1’ overlapped with juveniles caught in the Garnjarga area of the Teno mainstem (TEMS), while ‘Subpopulation 2’ overlapped juveniles caught 150km upstream in Inarijoki (INAR). Adults not assigned to either subpopulation cluster overlapped juveniles collected from other Teno locations, including the OUTA site midway between INAR and TEMS. Thus, the subpopulation structuring observed in the Teno mainstem by Aykanat et al. (2015) was generated by the sampling of migratory adults originating from two subpopulations with geographically distinct spawning and/or juvenile rearing sites.

**Figure 2:**
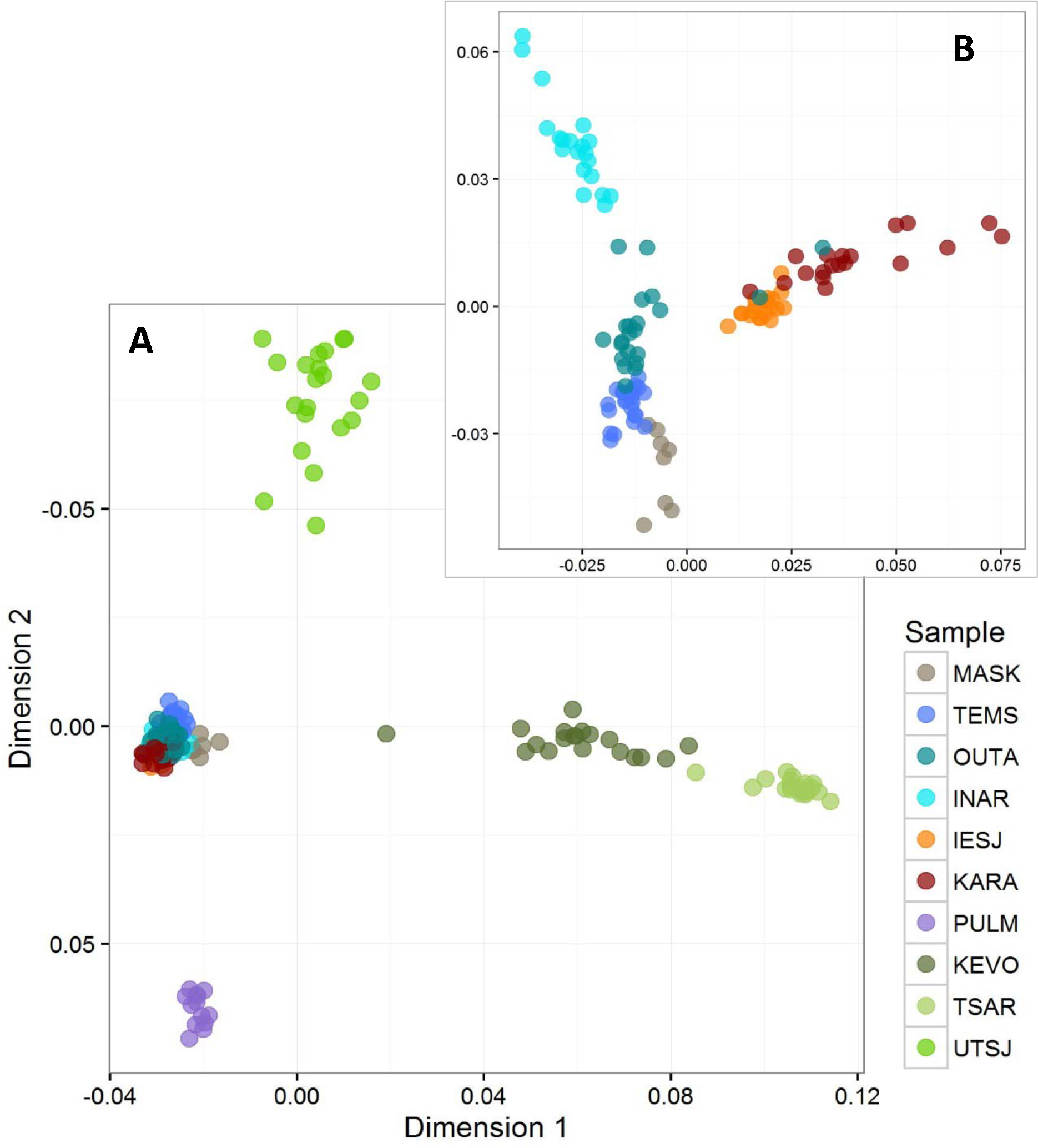
Two-dimensional MDS plot based on genome-wide IBS. (A) All juvenile samples; (B) all samples except KTVO, PULM, TSAR and UTSJ.

### Regions of the genome potentially responding to local selection

Distributions of test scores for the top-ranked 10,000 SNPs, and the top 0.5% that were retained, are shown in Fig. S4. By combining results from the outlier and environmental association tests, we identified 1,409 SNPs that, under our criteria, were candidates to be linked to variants under local selection (0.71% of total SNPs examined; 676 ‘OUT’ SNPs, 227 ‘PC1’ SNPs, 460 ‘PC2’ SNPs, Table S3). These ‘candidate SNPs’ occurred within 654 different haploblocks, and were associated with 522 overlapping and 258 downstream genes (Table S4; 1.9% of all genes annotated on the *S. salar* chromosomes; OUT: 258 overlapping/139 downstream; PC1: 93 overlapping/41 downstream; PC2: 168 overlapping/80 downstream). An additional 46 SNPs were supported as candidates by both XP-EHH and HapFLK results but no other tests (‘HH’ SNPs), however these were associated with only one additional gene (Tables S3 & S4). Thirty nine haploblocks (5%) contained SNPs on the 7K array that were identified as candidates for divergent selection or associated with traits of interest in previous studies (Table S3).)

Thirty-seven haploblocks had evidence for local selection both from outlier and/or environmental association analyses and from haplotype frequency/homozygosity analyses (Table 2; Fig. S5; 25 supported by HapFLK, 8 supported by XPEHH; 4 supported by both). These ‘candidate haploblocks’ were distributed over 17 of the 29 *S. salar* chromosomes. Three of the candidate haploblocks on Ssa09 (Fig. S5.5), five on Ssa12 (Fig. S5.8), two on Ssa17 (Fig. S5.15) and two on Ssa18 (Fig. S5.16) were located closely together on the chromosome and supported by a HapFLK signal that could be driven by a single selected locus; thus we considered there to be 28 independent candidate regions. Haploblock size ranged from 1.8Kb to 2.4Mb, reflecting substantial variation in local recombination rates around the candidate loci.

**Table 2:**
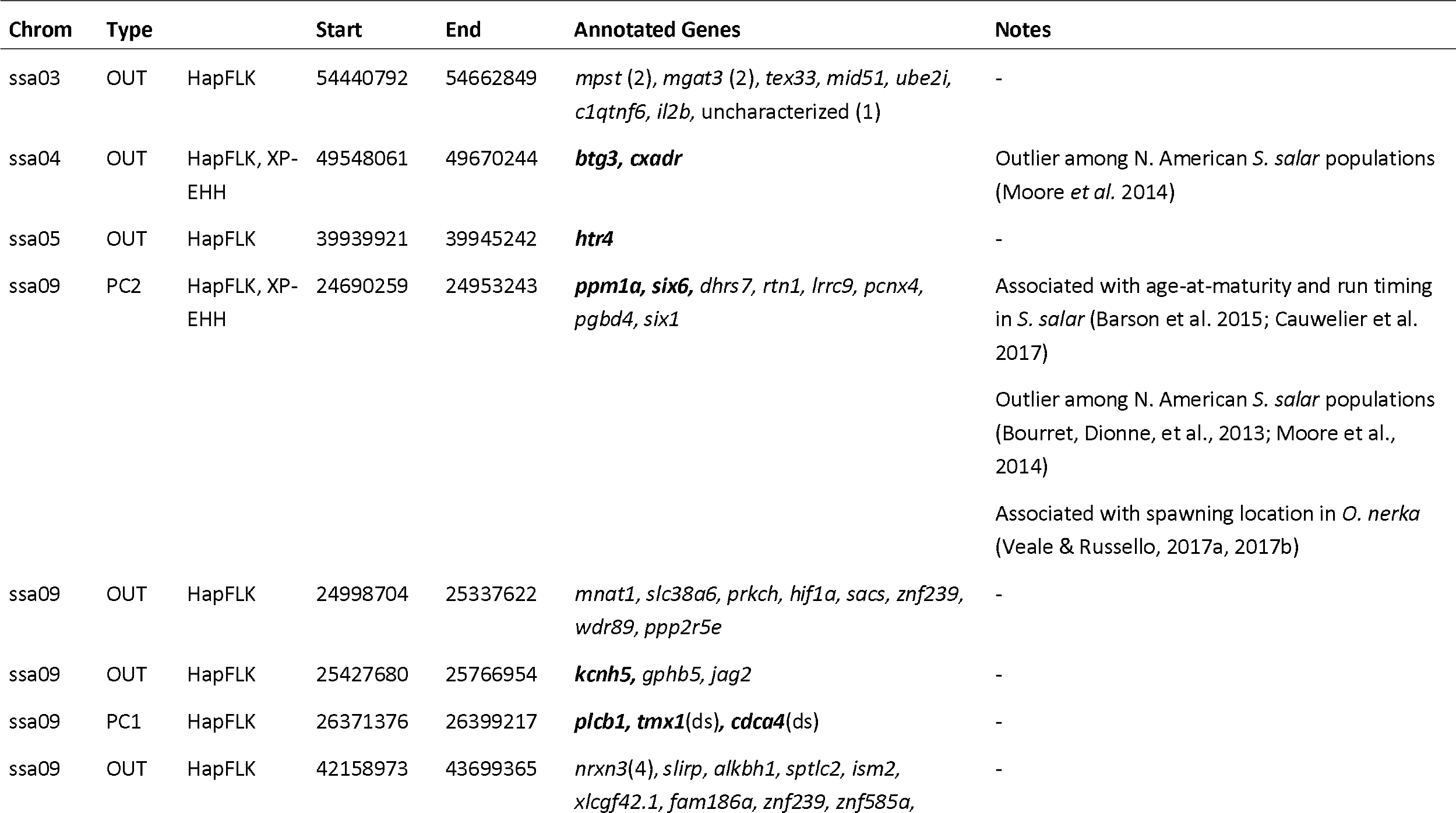
Candidate haploblocks supported by both outlier or environmental association analyses and HapFLK or XP-EIIII analyses. ‘Type’ – haploblock associated with PCI, PC2, or outlying (OU1), based on haploblock consensus SNP signal; ‘Annotated Genes’: gene symbols based on NCBI annotated gene products. Overlapping and selected downstream (ds) genes are shown. Genes in bold are those that are closest to the most likely selective targets (if any) based on visual observation of SNP score distribution (Fig. S5).

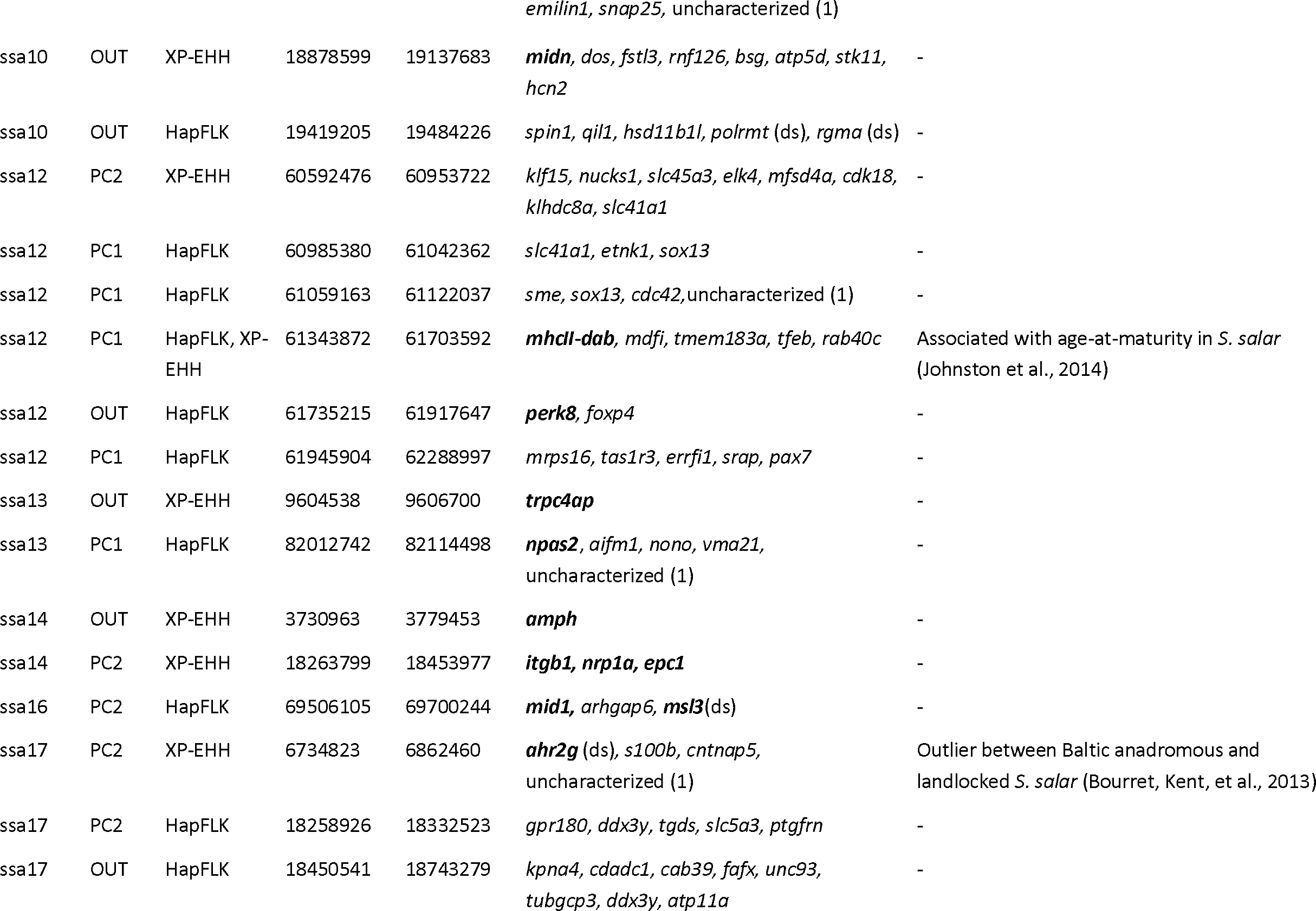

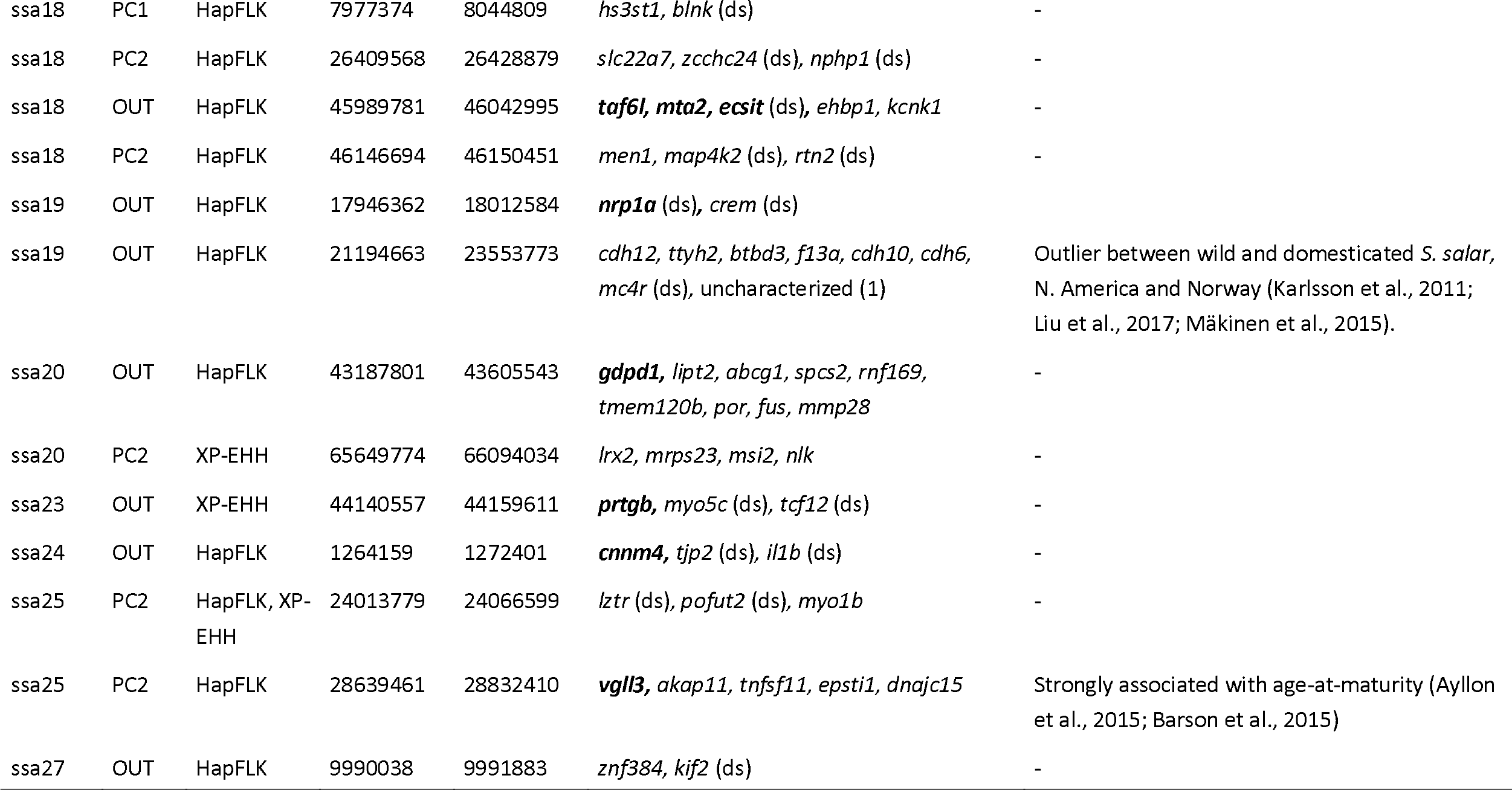

Six of these candidate haploblocks included loci of interest from previous studies (Table 2). A single haploblock with particularly strong evidence for divergent selection was located on Ssa09. *O. nerka* ecotype-associated RAD-tag 68810 mapped unambiguously to the *S. salar lrrc9* gene within this haploblock. Ecotype-associated RAD-tag 24343 did not align with the *S. salar* genome, but mapped unambiguously on the *O. mykiss* genome ≈ 36kb from 68810 between the genes *lrr9* and *dhrs7*. The zebrafish retinal *six6* enhancer mapped unambiguously 5’ of the *six6* gene within the haploblock.

### GO enrichment analysis

Extended GO term enrichment results are provided in Table S5. We restrict our discussion to GO enrichments significant at p < 0.001 (Table 3). We emphasize that GO term enrichment results, in general, can be skewed by the presence of a small number of well-studied genes in the dataset, and that gene functions in non-model organisms such as salmonids are not well characterized. Nevertheless, several general trends emerge from the results. Corresponding to expectations that adaptive divergence is likely to be mediated by changes in gene expression our total set of candidate loci is enriched for transcription factors (GO:0006367; GO:0035326). We find an over-representation of various neuronal components (including the those of the neuronal cell body, growth cone, and dendritic shaft), and a corresponding over-representation of processes involved in nervous system development and function, particularly in respect to the sensory system. We also find over-representation of cell surface/plasma member proteins, and corresponding enrichment of GO terms related to cell-cell adhesion. Finally, we observe a few highly enriched GO terms related to muscle and skeletal differentiation.

**Table 3:**
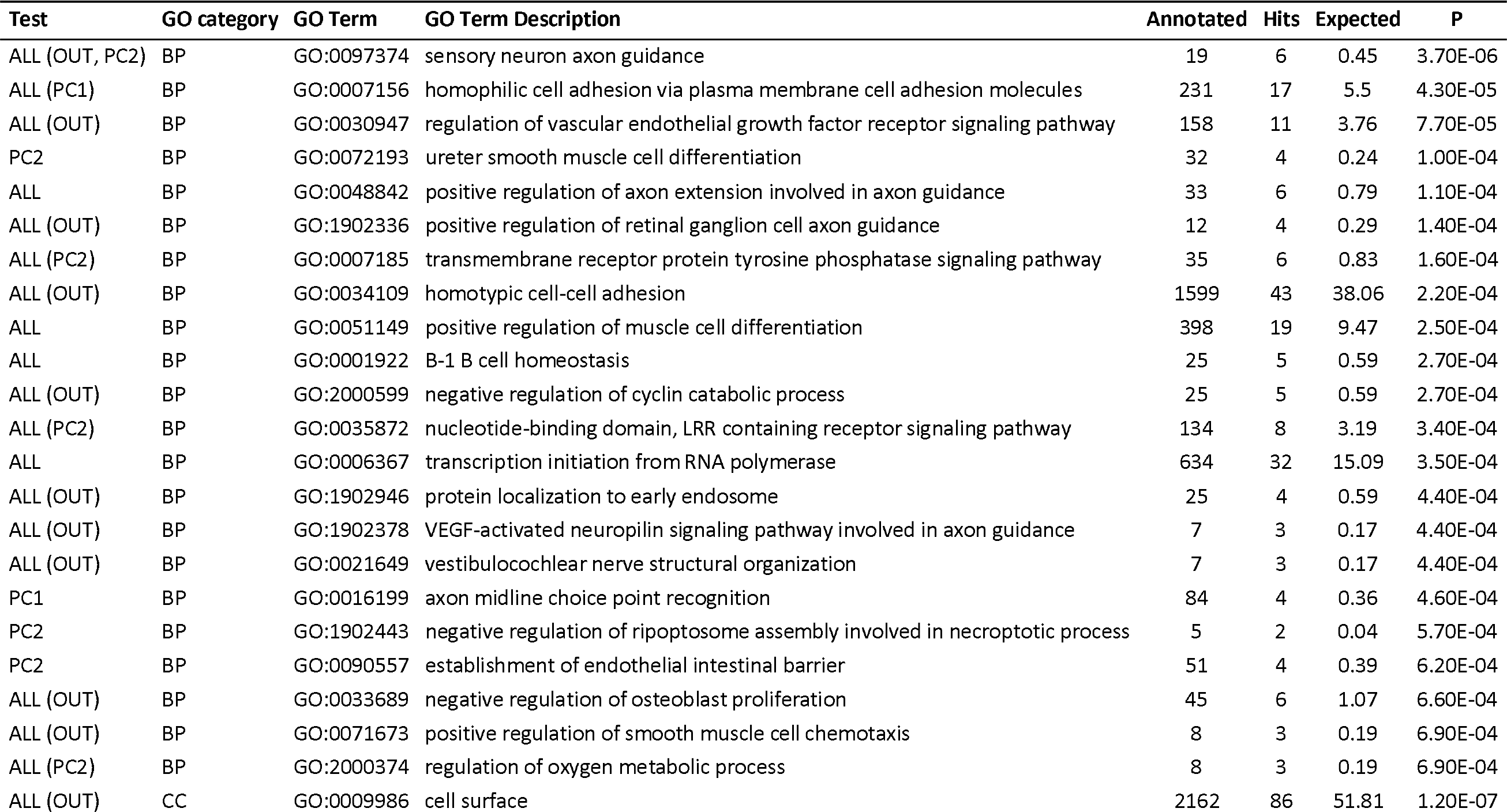
GO Enrichment Analysis. Only terms enriched at p < 0.001 are shown. Where GO terms are enriched under these criteria in more than one analysis, statistics show results from the largest set of examined genes, and additional analyses are shown in brackets. ALL: All candidate genes; PCI: candidate genes associated with PCI SNPs; PC2: candidate genes associated with PC2 SNPs; OUT: candidate genes associated with OUT SNPS; BP: Biological Process; CC: Cellular Component; ML: Molecular Function.

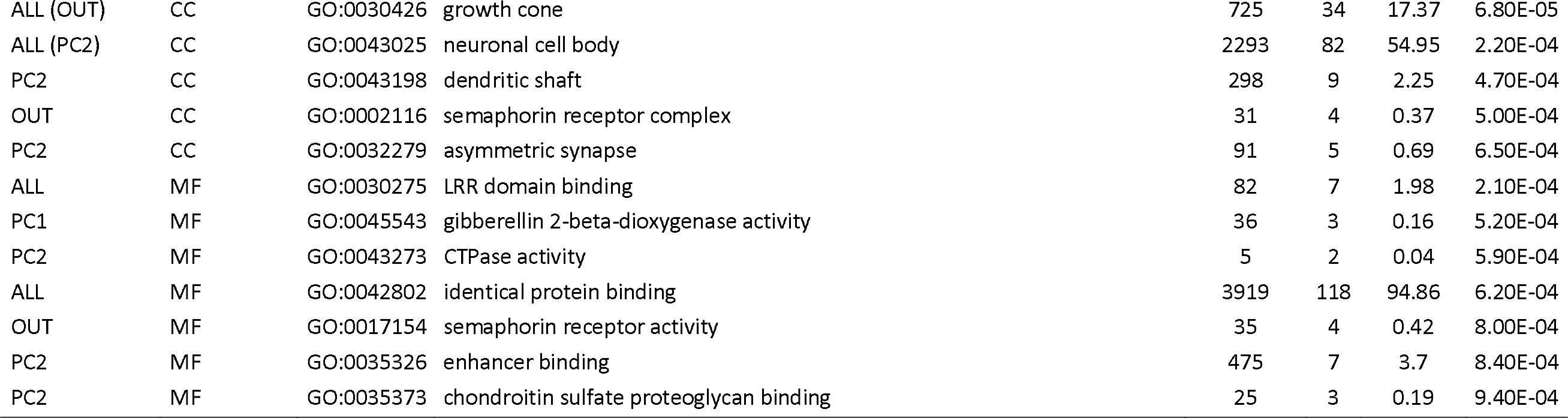

## DISCUSSION

In this study, we used a ‘bottom up’ approach – examining patterns of variation across the genome - to identify loci that may underlie fine-scale local adaptation of Atlantic salmon in a large river system. By combining results from outlier and environmental association analyses, we identified 1,409 candidate SNPs, annotated with 780 protein coding loci. While the relatively relaxed empirical p threshold (p<0.005 based on the entire distribution) means that these candidate genes may include numerous false positives, this threshold also increases our power to detect genes underlying polygenic adaptive traits (De Villemereuil, Frichot, Bazin, François, & Gaggiotti, 2014). By adding evidence from patterns of haplotype homozygosity, we found 28 regions of the genome that are particularly likely to contain loci under differential selection (Table 2, Fig. S5). Six of these regions have been documented as potential selective targets among other Atlantic salmon populations and/or contain candidate genes known to underlie ecologically relevant phenotypic variation (Table 2, Fig. S5). This observation both increases our confidence that the other regions identified using the same criteria also harbour genes under divergent selection, and also suggests that certain loci are under repeated selection among Atlantic salmon at local, regional and continental scales. Of particular note, we found a single genomic region that was highly differentiated among Teno River subpopulations, a candidate selective target throughout the range of Atlantic salmon, and which co-localized with an ecologically important haplotype within a different salmonid genus.

### A large-effect gene underlying age-at-maturity appears under differential selection among populations

A candidate haploblock for diversifying selection on Ssa25 contains the known large-effect locus underlying variation in age at maturity in Atlantic salmon (Ayllon et al., 2015; Barson et al., 2015; candidate gene *vgll3*) (Table 2, Fig. S5.26). Our results lend further support to the evidence in Barson et al. (2015) that variants at this locus may be differentially selected among rivers. This genomic region is strongly associated with PC2 (approximating flow volume), an association that suggests different trade-offs between size at maturity and reproductive success in different size rivers. At the simplest level, mechanical constraints could limit the access of larger, later-maturing fish to smaller tributaries, while only larger females may be able to successfully construct redds in higher-flow locations with coarser substrate (Kondolf & Wolman, 1993).

### A strong signature of divergent selection among Atlantic salmon populations co-localizes with a locus associated with spawning site selection in Sockeye salmon

We observe an extremely strong signal of diversifying selection approximately 2.5Mb along Atlantic salmon chromosome Ssa09 (Fig. 3, Table 2, Fig. S5.5). SNPs in this region are among the most extreme outliers in all three outlier analyses, are robustly associated with PC2 (approximating flow volume) in both environmental association analyses, and are clearly indicated as co-locating with a selective sweep by XP-EHH and HapFLK results. The selective signal centres on a cluster of closely linked SNPs spanning ≈7kB between the genes *protein phosphatase 1a (ppm1a)* and *SIX homeobox 6 (six6)*. The TEMS sample, from the large Teno mainstem, is almost fixed for one allele across these SNPs, and XP-EHH results indicate a selective sweep on this allele. Conversely, the samples from the smaller tributaries KEVO, TSAR and PULM are almost fixed for the alternate allele (Table S3). The strong relationship of allele frequencies at this location with PC2 infers that this outlying locus is also strongly associated with subpopulation run time, a relationship that we confirmed by performing a supplementary LFMM analysis with median run time as the dependent variable (Table S6).

**Figure 3:**
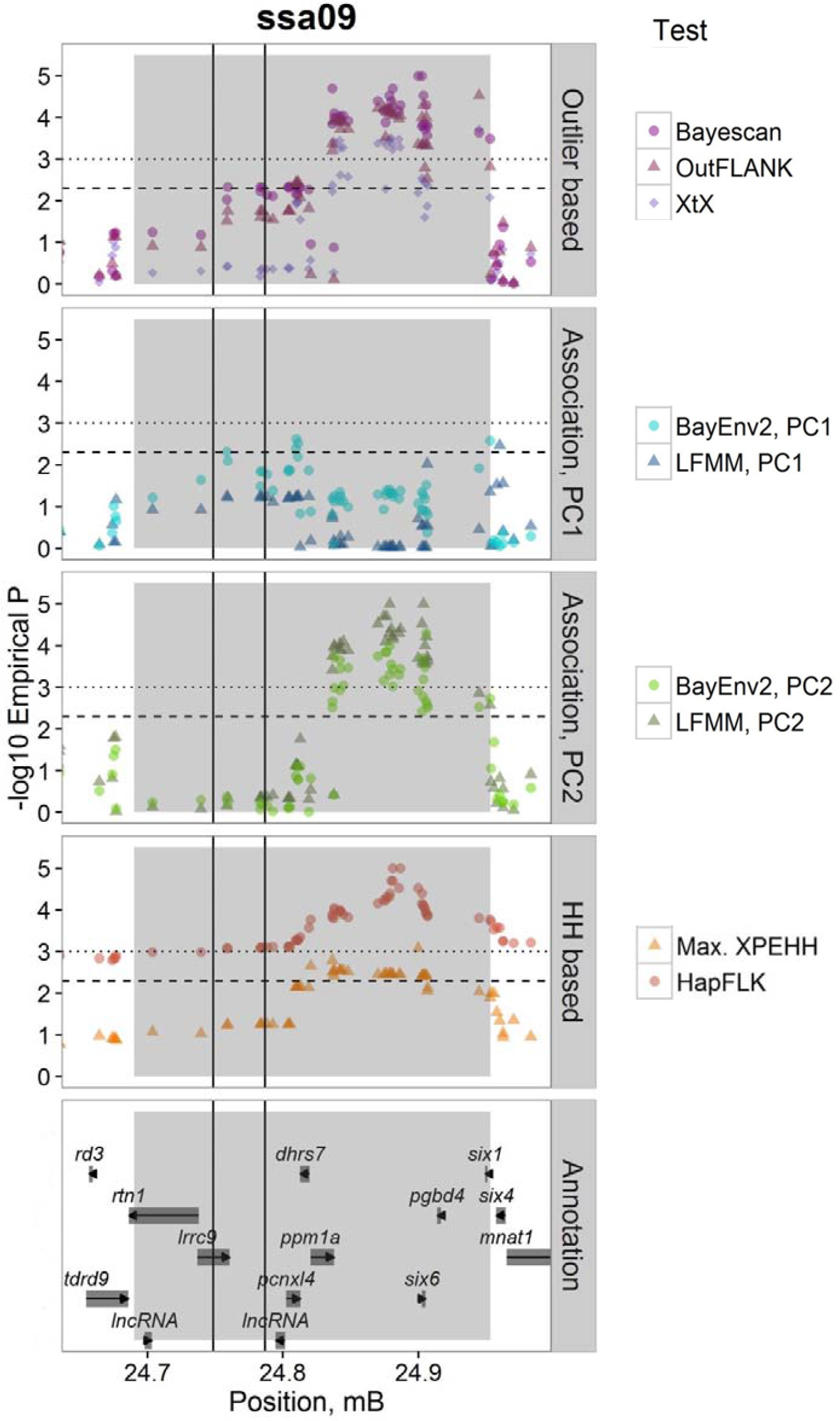
Signatures of local selection on the Ssa09 region containing the *Six6* gene. Dashed and dotted lines indicate empirical p < 0.005 and p < 0.001 respectively. The grey rectangle shows the boundaries of the associated haploblock. Black vertical lines indicate the mapping position *O. nerka* ecotype-associated RAD tags 68810 (left) and 24343 (right) (Veale & Russello, 2017b).

Differentiation of this Ssa09 region among Atlantic salmon populations was previously shown by Barson et al. (2015), where the candidate SNPs were F_st_ outliers and strongly associated with variation in age-at-maturity in genome-wide association studies before correction for population stratification. This pattern was observed both in the Teno and throughout Norway. Barson et al. (2015) also found a small but significant effect of genotype at this locus on length of returning adults. Recently, Cauwelier, Gilbey, Sampayo, Stradmeyer, & Middlemas (2017) found that the locus was associated with intra-population variation in run timing of Atlantic salmon in Scotland. Further, Bourret, Dionne, et al. (2013) and Moore et al. (2014), using a 7K SNP chip, found evidence that this region of Ssa09 was under diversifying selection among Atlantic salmon populations in North America. This suggests that the same variant may be involved in adaptive divergence throughout the range of the species. Intriguingly, this strongly outlying region is close to the locus associated with spawning site selection in *O. nerka* (Veale & Russello, 2017a). The relevant RAD-tags map within our candidate haploblock but ≈ 75 - 100Kb away from our strongest candidate markers (Fig. 3). Given the much lower density of the Veale & Russello (2017b), Nichols, Kozfkay, & Narum (2016), and Larson et al. (2017) RAD-tags compared to our SNPs, the same locus may be involved in local adaptation in both cases. Supporting this, SNPs on the 7K array found to be associated with age-at-maturity in Teno salmon by Johnston et al. (2014) map ≈700Kb distant (Table S3) but are clearly labelling our candidate locus. Further, Veale & Russello (2017a), sequencing through *lrrc9*, found that haplotype divergence increased towards our candidate region.

One gene flanking this region, *ppm1a*, is a broad-specificity enzyme whose potential role in mediating local adaptation is unclear. In contrast *six6* is an evolutionarily conserved transcriptional co-regulator with well characterized roles in development of the eye and establishment of the pituitary-hypothalamic axis in multiple vertebrates (Jean, Bernier, & Gruss, 1999; Seo, Drivenes, Ellingsen, & Fjose, 1998; Toy, Yang, Leppert, & Sundin, 1998). It is a paralogue of the invertebrate gene *optix*, which underlies locally adaptive variation in wing pigmentation across multiple butterfly genera (Zhang, Mazo-Vargas, & Reed, 2017). In the developing vertebrate eye, *six6* interacts with other genes to mediate both early initiation of the eye field and later establishment of the mature retina (Conte et al., 2010; Seo et al., 1998). It continues to be expressed in the fully developed eye (Aijaz et al., 2005), and is associated with thickness of the retinal nerve fibre layer and primary open-angle glaucoma in humans (Ulmer Carnes et al., 2014). Through its role in hypothalamic development, it is required for proper development of the suprachiasmatic nucleus, which is the central regulator of circadian timing in mammals (Clark et al., 2013), and may have a similar role in fish (Watanabe et al., 2012). *Six6* is also an important regulator of fertility in mammals of both sexes, via its effect on gonadotropin-releasing hormone production by the hypothalamus (Larder, Clark, Miller, & Mellon, 2011). Correspondingly, like *vgll3*, it is implicated in human pubertal timing (Hou et al., 2017). Barson et al. (2015) did not find any nonsynonymous mutations in the *six6* coding region, and results indicate that the selective target could be a regulatory element for this gene. Studies in various species have identified multiple 5’ evolutionarily conserved enhancer elements which regulate *six6* expression in different tissues and at different development stages (Conte et al., 2010; Ledford et al., 2017; Lee et al., 2012). *Six6* regulation also involves at least one 3’ enhancer (Conte et al., 2010; Ledford et al., 2017), and may also be mediated by local chromatin structure (Gómez-Marín et al., 2015). We found that one conserved 5’ enhancer, responsible for *six6* expression in the maturing medaka retina (Conte et al., 2010) maps between two candidate SNPs (ss1868043109 and ss1868793364). Barson et al. (2015) showed that a second conserved enhancer, necessary for *six6* expression in the mouse forebrain (Lee et al., 2012) maps close to candidate SNP ss1868315338. Additional putative enhancers, with poorly characterized functions, are also located within our candidate region in other species (Conte et al., 2010; Ledford et al., 2017; Lee et al., 2012).

How this region interacts with the Ssa25 (*vgll3*) locus to influence age-at-maturity – whether through a direct functional relationship, and/or indirectly by altering the selective landscape for the Ssa25 locus among populations – remains unknown Given the implication that the same genomic region may be involved in spawning site differentiation in *O. nerka*, we hypothesize that *six6* could mediate local adaptation/spawning site differentiation in both species by modulating aspects of the sensory and/or reproductive system, including reproductive timing. Notably, both Atlantic salmon in larger rivers and sockeye salmon on lake shores tend to be spawning at increased depths (with consequent altered light regimens) and on coarser substrates (Frazer & Russello, 2013; Louhi, Mäki-Petäys, & Erkinaro, 2008). Further, *O. nerka* spawning ecotypes also differ in their reproductive timing (Frazer & Russello, 2013). As with its invertebrate paralogue (Zhang et al., 2017), *six6* may have pleiotropic effects on multiple aspects of the phenotype by acting as ‘master switch’ across different regulatory networks during development.

In addition to *vgll3* and *six6*, several other genes implicated in human pubertal timing (Hou et al., 2017) are associated with PC2 SNPs. These include *neuronal growth regulator 1* (*negr1*, Ssa10), linked to both obesity and sexual maturation in several taxa (Lee et al., 2012); *trmt11* (Ssa05); *ptprf* (Ssa10); *ntrk2* (Ssa10); *lgr4* (Ssa11); *ptprd* (Ssa18); *kcnk9* (Ssa27); and *h6st1* (Ssa29) (Table S4). These genes merit further investigation as potential interactors involved in the relationship between *six6*, *vgll3* and life-history variation among different Atlantic salmon populations.

### Possible selection on genes involved in circadian timing

After the Ssa09 locus, the genomic region most clearly associated with PC2 (and run timing) in the LFMM analysis is located ≈ 19Mb along Ssa11 (Table S3, Table S4, Table S6).

While this region is not supported as a ‘candidate haploblock’ given our current thresholds, it contains SNPs within the top 1% of XP-EHH scores. This region is also highly differentiated between northern and southern populations of *S. salar* in Norway (Kjærner-Semb et al., 2016). One of two candidate genes at this locus is zinc finger homeobox 3 (*zfhx3*), a transcription factor expressed in the suprachiasmatic nucleus with a role in mammalian circadian rhythms, including sleep (Balzani et al., 2016). This must therefore be considered a candidate locus for co-selection with *six6*.

A candidate haploblock at on Ssa13 overlaps *neuronal PAS domain protein 2* (*npas2*) (Table 2, Fig. S5.10). *Npas2* is a paralogue of *clock*, which is well known for regulating circadian rhythms and reproductive cycles in diverse taxa. In mammals, the two genes have similar roles in the suprachiasmatic nucleus, and *npas2* can compensate if *clock* is silenced (DeBruyne, Weaver, & Reppert, 2007). *Clock* polymorphisms are associated with the timing of reproduction in rainbow trout and chinook salmon, and show a latitudinal cline consistent with local selection in the latter (Leder, Danzmann, & Ferguson, 2006; O’Malley, Camara, & Banks, 2007; O’Malley & Banks, 2008). Few salmonid studies have examined *npas2*, but O’Malley, Jacobson, Kurth, Dill, & Banks (2013) found that *npas2*-linked variation discriminated O. *tshawytscha* populations with different migratory timing. In our analysis, however, this candidate haploblock is not associated with PC2, meaning that there is no clear link with run time.

### Evidence for local selection on genes mediating energy homeostasis

The extremely large candidate haploblock on Ssa19 (Table 2, Fig. S5.20) overlaps a region of the genome responding to domestication selection in North America *S. salar* (Liu et al., 2017; Mäkinen et al., 2015). This region also contains a SNP differentiating wild and domestic Norwegian *S. salar* (Karlsson et al., 2011). Following Liu et al. (2017), we hypothesize that the target of selection is a regulatory element for the downstream *melanocortin receptor 4* (*mc4r*) gene. This gene is a controller of energy homeostasis and somatic growth in fish and other vertebrates via its influence on food intake and energy expenditure (Krashes, Lowell, & Garfield, 2016; Metz, Peters, & Flik, 2006); it is a well-characterized human obesity gene and expected to be differentially selected in the wild vs. domestic environment. Several other obesity-associated genes are candidate selective targets: four copies of *neurexin 3* (*nrxn3*, Heard-Costa et al. 2009) on a large candidate haploblock on Ssa09 (Table 2, Fig. S5.6); *negr1* (discussed above); *lingo2* (Ssa05 and Ssa09); *arid5b* (Ssa01); and *kcdt15* (Ssa11) (Claussnitzer et al., 2015; Castillo, Hazlett & Orlando, 2017; Table S4). The observation of these genes as selective candidates conforms to a model in which different Teno Atlantic salmon sub-populations are adapted to different energy-balance optima. Further, as threshold levels of fat reserves at specific times of the year are thought influence sexual maturity in salmon (Thorpe, Mangel, Metcalfe, & Huntingford, 1998), such obesity-associated genes may further interact with *vgll3* and *six6* to influence variation in life-history dynamics among Atlantic salmon populations.

### Signatures of directional selection on MHCII and other genes involved in pathogen response

A clear signal consistent with directional selection occurs around the *major histocompatibility complex II* (*mhcII*) gene on Ssa12 (Table 2, Fig. S5.8). Allelic variation within this haploblock correlates with PC1 (approximating distance from the ocean). Salmonid genomes contain only a single classical *mhcII* locus (Gómez, Conejeros, Marshall, & Consuegra, 2010), which mediates an immune response to bacteria and parasites (Piertney & Oliver, 2006). Elevated *mhc* diversity confers broader pathogen resistance: thus, over evolutionary timescales, *mhc* alleles are maintained by balancing selection (Piertney & Oliver, 2006). However, pathogen pressure within a population can generate directional selection on particular *mhc* alleles over shorter time scales. In salmonids, specific *mhcII* alleles have been associated with resistance to the bacterial diseases piscirickettsiosis (Gómez, Conejeros, Consuegra, & Marshall, 2011), furunculosis (Kjøglum, Larsen, Bakke, & Grimholt, 2008; Langefors, Lohm, Grahn, Andersen, & von Schantz, 2001) and infectious salmon anaemia (Kjøglum, Larsen, Bakke, & Grimholt, 2006), and the parasite *Myxobolus cerebralis* (Dionne, Miller, Dodson, & Bernatchez, 2009). Similar evidence for localized directional selection on *mhcII* variants has been observed in sockeye salmon at both broad and fine spatial scales (Larson, Seeb, Dann, Schindler, & Seeb, 2014; McClelland et al., 2013).

Signatures of diversifying selection around other genes involved in pathogen response supports this model of different pathogen pressures among our sampled subpopulations, despite their co-occurrence in the same drainage basin. A strong XP-EHH signal suggesting a recent selective sweep, in particular in the Yla Pulmankijarvi (PULM) subpopulation, occurs on Ssa04 around a copy of *coxsackie virus and adenovirus receptor* (*cxadr*) (Table 2, Fig. S5.2). Although this gene has multiple functions, this may indicate virus-mediated selection. Other immune associated genes located in candidate haploblocks include two important mediators of the inflammatory response: *beta 1 integrin* (*itgb1*), on Ssa14 and *interleukin 1 beta* (*il1b*) on Ssa24 (Table 2).

### Other candidate selective targets

The candidate haploblock on Ssa17, associated with PC2, is immediately upstream of *aryl hydrocarbon receptor 2 gamma* (*ahr2g*) (Table 2, Fig. S5), and we observed candidate SNPs upstream of two of the three other ahr2 genes annotated on the *S. salar* genome (*ahr2a* and *ahr2d*, Table S4). Although genetic variation in the aryl hydrocarbon receptor pathway underlies adaptation to anthropogenic pollution (Reid et al., 2016; Wirgin et al., 2011), this is unlikely to be a selective force in the Teno. Regulatory variation in the Ahr2 pathway is associated with ecotypic diversity in craniofacial morphology in Arctic charr (*Salvelinus alpinus*, Ahi et al., 2014).

The small candidate haploblock on Ssa05 overlaps one of three conserved copies of *5-hydroxytryptamine receptor 4* (*htr4*) annotated on the Atlantic salmon genome (Table 3, Fig. S5). The neurotransmitter 5-HT (serotonin) is an important modulator of vertebrate behavioural state; in Atlantic salmon, for example, it mediate the expression of different behavioural syndromes between early and later-emerging fry, likely an adaptive response to varying levels of competition (Thornqvist, Hoglund, & Winberg, 2015). The class 4 receptors of 5-HT are not well characterized in fish, however in mammals they are expressed in the nervous and gastrointestinal systems, and have specific roles in memory formation and feeding behaviour (Bockaert, Claeysen, Compan, & Dumuis, 2008).

Examination of GO terms implied an enrichment of nervous system genes overlapping or downstream of candidate SNPs (Table 3). Further, a number of candidate haploblocks and SNPs are annotated with genes whose adaptive function is not clear but which code proteins important for neuronal function. These include *amphiphysin* (*amph*), the only gene annotated on the candidate haploblock near the start of Ssa14 (Table 2; Fig. S5.11), and two copies of *neurophilin 1a* (*nrp1a*) associated with Ssa14 and Ssa19 haploblocks (Table 2; Figs. S5.12 & S5.19). We tentatively hypothesize that aspects of the nervous system, perhaps underlying sensory systems involved in natal homing, may be under differential selection among different Atlantic salmon populations. This suggests several interesting avenues for future research.

### Management Implications

Our results strongly indicate that Teno River Atlantic salmon subpopulations identified on the basis of microsatellite variation are also differentiated at the functional genetic level, and so are unlikely to be ecologically interchangeable. This underscores the recommendations of Vähä, Erkinaro, Niemelä, & Primmer (2007) and Vähä, Erkinaro, Falkegård, Orell, & Niemelä (2017) that the stock be managed at the subpopulation level. On a broader scale, our results support the hypothesis that local adaptation is common among Atlantic salmon populations, even those in geographic proximity. This should be taken into account, for example, in stocking and restoration programs.

## ACKNOWLEDGEMENTS

This work was supported by Academy of Finland Grants 284941 and 314254 to CRP and a grant from the Finnish Cultural Foundation to VLP. We thank Sigbjørn Lien, Matthew Kent, and Silje Karoliussen (CIGENE) for SNP Array genotyping, Kristiina Haapanen for laboratory assistance, and Matthieu Bruneaux and Ksenia Zueva for help with GO analyses. Computing resources were provided by CSC – IT Center for Science Ltd (Finland).

## AUTHOR CONTRIBUTIONS

CRP, HM & VLP designed the study. JE,J-PV & PO collected tissue samples. HM &J-PV performed laboratory work. VLP analysed the data and wrote the manuscript, with input from CRP, HM, JE & PO. All authors approved the manuscript.

## DATA ACCESSIBILITY

SNP genotypes, raw analysis results, code and other relevant files will be deposited in the Dryad Digital Repository.

---

**Table S1:** Results of Principle Component Analysis.

| Rotation: |  |  |  |  |
| --- | --- | --- | --- | --- |
|  | PC1 | PC2 | PC3 | PC4 |
| Catchment | 0.191 | 0.957 | 0.199 | 0.091 |
| Distance | -0.582 | 0.266 | -0.427 | -0.639 |
| Elevation | -0.632 | 0.095 | -0.192 | 0.744 |
| Accessibility | -0.474 | -0.068 | 0.861 | -0.172 |

| Importance: |  |  |  |  |
| --- | --- | --- | --- | --- |
|  | PC1 | PC2 | PC3 | PC4 |
| Standard_deviation | 1.522 | 0.986 | 0.799 | 0.274 |
| Proportion_variance | 0.579 | 0.243 | 0.160 | 0.019 |
| Cumulative_prop | 0.579 | 0.822 | 0.981 | 1.000 |

**Table S2:** Pairwise unbiased F_st_ among samples, based on all SNPs.

|  | IESJ | INARI | KARA | KEVO | MASK | OUTA | PULM | TEMS | TSAR | UTSJ |
| --- | --- | --- | --- | --- | --- | --- | --- | --- | --- | --- |
| IESJ | 0.000 | 0.024 | 0.016 | 0.068 | 0.035 | 0.013 | 0.084 | 0.017 | 0.129 | 0.062 |
| INARI |  | 0.000 | 0.028 | 0.075 | 0.046 | 0.016 | 0.095 | 0.024 | 0.136 | 0.071 |
| KARA |  |  | 0.000 | 0.075 | 0.044 | 0.018 | 0.092 | 0.023 | 0.136 | 0.069 |
| KEVO |  |  |  | 0.000 | 0.083 | 0.064 | 0.128 | 0.065 | 0.059 | 0.084 |
| MASK |  |  |  |  | 0.000 | 0.033 | 0.104 | 0.033 | 0.148 | 0.080 |
| OUTA |  |  |  |  |  | 0.000 | 0.084 | 0.008 | 0.124 | 0.059 |
| PULM |  |  |  |  |  |  | 0.000 | 0.084 | 0.188 | 0.124 |
| TEMS |  |  |  |  |  |  |  | 0.000 | 0.125 | 0.057 |
| TSAR |  |  |  |  |  |  |  |  | 0.000 | 0.140 |
| UTSJ |  |  |  |  |  |  |  |  |  | 0.000 |

**Table S3** (separate.csv file): Candidate SNPs. **SNP**: Candidate SNP ss#; **Chr**: Chromosome (ssa); **Position**: Position on chromosome (bp); **EmpP:** Empirical P values, based on: **Bayescan**: Empirical P value based on ?; **OutFLANK**:; **XtX:** BayEnv2 XtX score; **BayenvPC1**: BayEnv2 |p| - association with PC1; **BayenvPC2:** BayEnv2 |p| - association with PC2; **LFMMPC1:** LFMM z-score - association with PC1; **LFMMPC2:** LFMM z-score - association with PC2; **HapFLK**: HapFLK score; **XPEHH**: Maximum |XP_EHH| over nine interpopulation comparisons; **Type**: SNP classification, OUT, PC1, PC2, EHH; **XPEHH/HapFLK**: SNP is in the top 0.05% for either XP-EHH or HapFLK (1), or both (2); **IESJ_maf**; **INAR_maf**; **KARA_maf**; **KEVO_maf**; **MASK_maf**; **OUTA_maf**; **PULM_maf**; **TEMS_maf**; **TSAR_maf**; **UTSJ_maf**: Minor allele frequency in each sample (minor allele determined over all samples). **HB_start**; **HB_end**: Haploblock boundaries (bp); **7K_Outlier**: name of 7K SNP within haploblock; **7K_Study:** study in which 7K SNP was identified, (1) Bourret et al. 2013a; (2) Bourret et al. 2013b; (3) Moore et al. 2014; (4) Jeffery et al. 2017; (5) Mäkinen et al. 2015; (6) Liu et al. 2017; (7) Karlsson et al. 2011; (8) Gutierrez et al. 2016; (9) Johnston et al. 2014; (10) Perrier et al. 2013.

**Table S4** (separate.csv file): Candidate SNP annotation, overlapping or closest downstream genes. **SNP:** Candidate SNP ss#; **Chr**: Chromosome (ssa); **Position:** Position on chromosome (bp); **Type**: Candidate SNP classification (OUT, PC1, PC2, EHH); HapFLK/XPEHH: Number of HapFLK/XP-EHH tests in which SNP is in the top 0.5%; **Hit:** Overlapping or closest downstream gene; **Strand:** gene strand; **Distance:** gene distance from SNP; **Product:** gene product inferred by NCBI; **Gene_symbol**: gene symbol: **SNP_location:** SNP location in respect to gene; **Gene_Location:** gene location in respect to SNP; **Alt_Hit:** Closest gene, if not downstream; **Alt_Strand:** gene strand; **Alt_Distance:** gene distance from SNP; **Alt_Product:** gene product inferred by NCBI; **Alt_Symbol:** gene symbol; **Alt_Location:** gene location in respect to SNP; **Noncoding_hit:** closest or overlapping non-coding locus, if closer than coding loci; **NC_strand:** locus strand; **NC_distance:** locus distance from SNP; **NC_type:** locus type; **NC_Location:** locus location in respect to SNP.

**Table S5** (separate.csv file): Extended GO term analyses. All GO terms enriched at p>0.01 are reported. **Test**: whether result was for genes annotated on all SNPs, PC1 candidate SNPs, PC2 candidate SNPs, or OUT candidate SNPs; **Aspect:** BP, Biological Process; CC, Cellular Component; MF, Molecular Function; **Annotated:** number of background genes annotated with the GO term; **Significant:** number of candidate genes annotated with the GO term; **Expected:** expected number of candidate genes with the GO term; **classicFisher:** uncorrected significance of GO term enrichment.

**Table S6:**
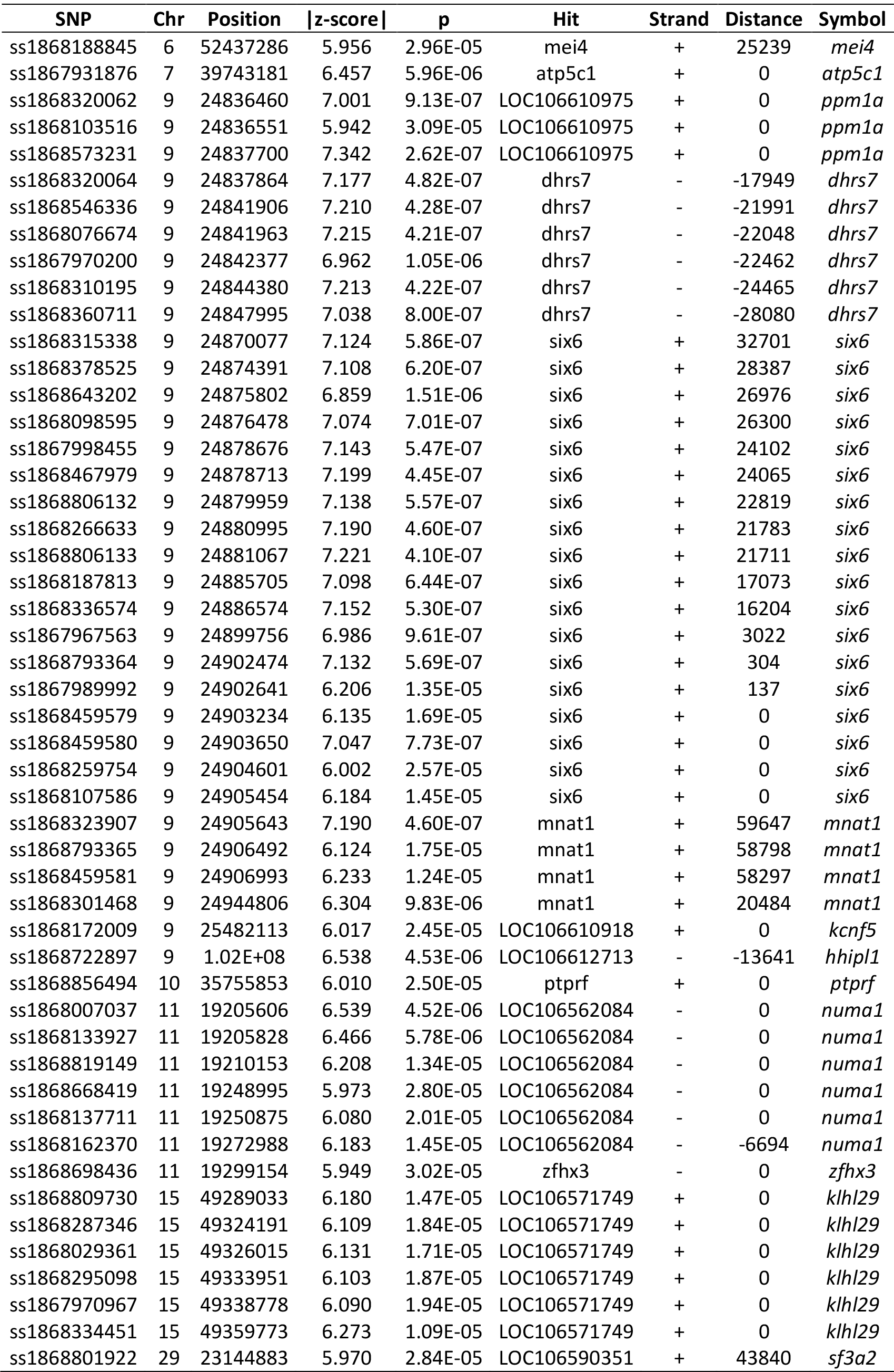
Top 50 SNPs associated with population run-time in the supplementary LFMM analysis. SNPs are ordered by genomic position. **SNP**: dbSNP ss#; **Chr**: Chromosome; **Position**: Position (bp); **|z-score|**: Absolute z-score, median over five runs; **p**: Probability adjusted for λ; **Hit**: overlapping or closest downstream gene; **Strand**: gene strand; **Distance**: distance of gene from SNP; **Symbol**: gene symbol.

